# Radioprotective drug screening in a salivary gland tissue chip

**DOI:** 10.1101/2023.02.06.527345

**Authors:** L. Piraino, C.Y. Chen, J. Mereness, P. M. Dunman, C. E. Ovitt, D. S. W. Benoit, L. A. DeLouise

## Abstract

Ionizing radiation damage to the salivary glands during head and neck cancer treatment often causes a permanent loss of secretory function. Due to the resulting decrease in saliva production, patients experience difficulty with eating, speaking, and swallowing and are predisposed to oral infections and tooth decay. While the radioprotective drug amifostine is approved to prevent radiation-induced hyposalivation, it has intolerable side effects that limit its use and motivate research into discovering alternatives. To address this issue, we have developed a salivary gland mimetic (SGm) tissue chip platform for use in high-content drug discovery. Here, we report on the development and validation of in-chip assays to quantify reduced glutathione and cellular senescence (β-galactosidase) as measures of radiation damage and protection using WR-1065, the active form of amifostine. Following validation, we next tested our assays using other reported radioprotective drugs including Edaravone, Tempol, N-acetylcysteine, Rapamycin, Ex-Rad, and Palifermin. The validated assays were then used to screen a library of FDA-approved compounds for radioprotection. We screened 438 compounds, obtained 25 hits that were further tested for EC_50_ values and downselected using information from the PubChem database. Lead compounds were identified that are being tested in preclinical models.

## 1. Introduction

Ionizing radiation damage to the salivary glands during head and neck cancer treatment often causes a permanent loss of secretory function and reduced salivary flow. Due to decreased saliva production, patients experience difficulty with eating, speaking, and swallowing [1,2]. Additionally, patients are at an increased risk of oral infections and tooth decay and suffer a reduced quality of life [2,3]. Current treatment options, including sialogogues, mouthwashes, and chewing gum, only provide temporary relief and there is no cure [2]. Several strategies have been proposed to alleviate this damage, including cell/hydrogel transplantation [4–7] and gene therapy [8–11]. Despite promising results, these methods have remained experimental and are targeted toward patients already experiencing xerostomia. Hence, there is an unmet need to provide current and future head and neck cancer patients with preventative therapies to protect salivary gland function. Currently, intensity modulated radiation therapy (IMRT) and the radioprotective drug amifostine are used clinically to prevent salivary gland damage. IMRT involves using 3D imaging to target the radiation beams at the cancer and away from sensitive organs such as the salivary gland [12]. While this method can be beneficial in some cases, there are mixed results on patient-reported claims of dry mouth and it cannot be implemented in some cases due to tumor location [12,13]. Amifostine is the only FDA-approved drug to prevent radiation-induced xerostomia, but it is often discontinued during fractionated radiation regimens due to severe side effects, including nausea, vomiting, and hypotension [14,15]. Additionally, its short half-life *in vivo* limits its efficacy, as the drug is cleared within minutes of administration [16]. These drawbacks highlight the critical need to discover new radioprotective drugs to prevent xerostomia.

Our approach for addressing this issue is based on the development and use of a salivary gland tissue chip for high-content drug screening. Recently, we developed a microbubble (MB) array-based tissue chip consisting of 3D salivary gland tissue mimetics (SGm) cultured in a polyethylene glycol (PEG) hydrogel microenvironment that recapitulates physiological conditions [17]. The spherical architecture of the MB creates a distinct niche that promotes cell viability, expression of acinar cell proteins, and functional calcium signaling [17]. In addition, we validated the use of this platform for radioprotection studies by using immunohistochemical staining of individual SGm to enumerate foci of DNA damage markers γH2AX and 53BP1 after irradiation. Analysis of control chips versus chips treated with WR1065, the active form of amifostine, showed a reduction in DNA damage with drug treatment [17].

To enable high-content drug screening, array-based assays that simultaneously interrogate each SGm on the chip (~280 MBs/cm^2^) are necessary. Based on literature, several assays commonly used to report radiation-induced cellular damage were tested in the tissue chip format. Assays tested included reactive oxygen species (ROS) generation and mitigation, apoptosis, secretion, and cytotoxicity at various time points post-radiation (Table S5.1). Based on signal to noise ratio and reproducibility, a reduced glutathione assay [18] and a cellular senescence [19] assay were selected for further development for high-content screening of radioprotective drugs.

These assays were first tested with 0 Gy, 15 Gy, and 15 Gy + 4 mM WR-1065 to assess their ability to detect radiation damage and WR1065-mediated radioprotection of individual SGm within the MB-hydrogel tissue chip (40-50 MBs per chip). The assays were then tested on other known radioprotective drugs including Tempol [20,21], N-acetylcysteine [22], Edaravone [23], Rapamycin [24], Ex-Rad [25,26], and Palifermin [27,28] to evaluate their suitability for high-content measurement of radioprotection. A library of FDA-approved compounds was screened for radioprotection using the assays, identifying 25 hits with both assays (double hits). These double hits were further investigated for their suitability as potential therapeutics using database searches and experimental determination of dose-response relationships.

## 2. Materials and Methods

### Materials

Information for the drugs used for assay development and validation are shown in Table 1. Drug screening was completed using a library of FDA-approved compounds (Selleck Chemicals).

**Table 1:**
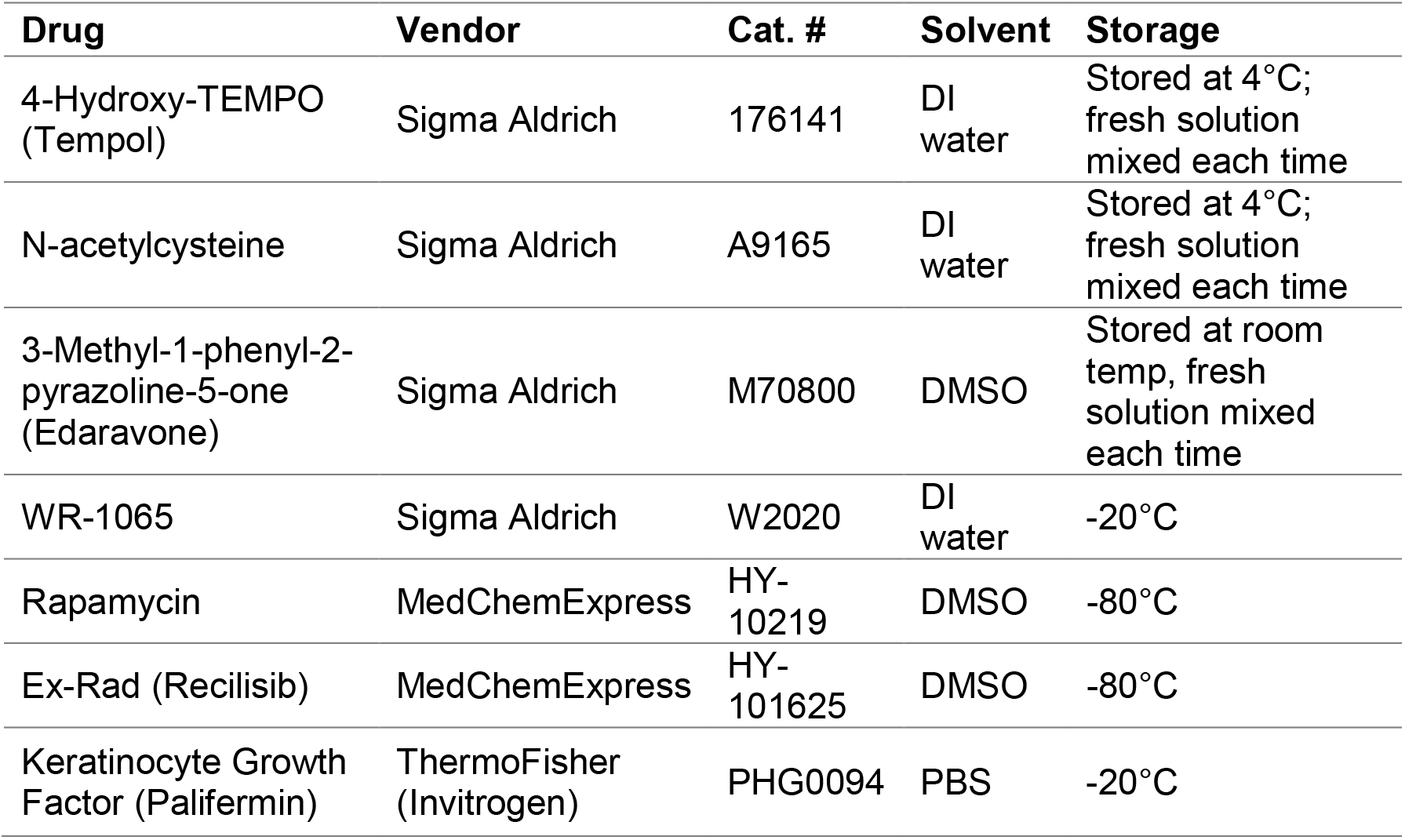
Product information for radioprotective drugs used in this study

| Drug | Vendor | Cat. # | Solvent | Storage |
| --- | --- | --- | --- | --- |
| 4-Hydroxy-TEMPO (Tempol) | Sigma Aldrich | 176141 | DI water | Stored at 4°C; fresh solution mixed each time |
| N-acetylcysteine | Sigma Aldrich | A9165 | DI water | Stored at 4°C; fresh solution mixed each time |
| 3-Methyl-1-phenyl-2-pyrazoline-5-one (Edaravone) | Sigma Aldrich | M70800 | DMSO | Stored at room temp, fresh solution mixed each time |
| WR-1065 | Sigma Aldrich | W2020 | DI water | -20°C |
| Rapamycin | MedChemExpress | HY-10219 | DMSO | -80°C |
| Ex-Rad (Recilisib) | MedChemExpress | HY-101625 | DMSO | -80°C |
| Keratinocyte Growth Factor (Palifermin) | ThermoFisher (Invitrogen) | PHG0094 | PBS | -20°C |

Lead compounds from the drug screen were purchased from Selleck Chemicals for dose-response studies (Table 2) and prepared and stored per manufacturer’s instructions.

**Table 2:** Compounds identified as double hits in the drug screen. Catalog numbers are from Selleck Chemicals.

| **Drug** | **Cat. #** |
| --- | --- |
| Adefovir Dipivoxil | S1718 |
| Albendazole | S1640 |
| Calcitriol | S1466 |
| Cilostazol | S1294 |
| Curcumin | S1848 |
| Didanosine | S1702 |
| Diethylstilbestrol | S1859 |
| Doripenem Hydrate | S1374 |
| Emtricitabine | S1704 |
| Enoxacin | S1756 |
| Eplerenone | S1707 |
| Etidronate | S1857 |
| Gemcitabine | S1714 |
| Glipizide | S1715 |
| Indomethacin | S1723 |
| Lomustine | S1840 |
| Melatonin | S1204 |
| Meropenem | S1381 |
| Nifedipine | S1808 |
| Oxcarbazepine | S1391 |
| Phenylbutazone | S1654 |
| Prazosin HCl | S1424 |
| Rifabutin | S1741 |
| Rifampin | S1764 |
| Triamcinolone | S1933 |

### Animals

Female SKH mice, backcrossed 6 generations on C57BL/6J mice, aged 6-12 weeks were used in this study. Only female mice were used due to known sex differences in rodent salivary glands, with female glands more accurately emulating human salivary gland structure and function [29,30]. Animals were maintained on a 12 hr light/dark cycle and group-housed with food and water available *ad libitum*. All procedures were approved and conducted in accordance with the University Committee on Animal Resources at the University of Rochester Medical Center (UCAR #2010-24E).

### Microbubble (MB) array fabrication

Microbubble (MB) arrays were fabricated in poly(dimethyl) siloxane (PDMS) using gas expansion molding as previously described [17,31,32]. PDMS (Dow Corning Sylgard 184) was mixed in a 10:1 ratio of base to curing agent and poured over a silicon wafer template consisting of deep etched cylindrical pits with a 200 μm diameter, spaced 600 μm apart. The PDMS was cured at 100 °C for 2 hrs before peeling off the template, resulting in an array of spherical cavities with 200 μm opening and ~350 μm diameter. Circular chips with 0.7 cm diameter (48 well) or 0.5 cm diameter (96 well) were punched from the casted PDMS and glued into well plates using a 5:1 ratio of PDMS. The MBs were primed in a desktop vacuum chamber with 70% ethanol to facilitate air removal from the MBs and replacement with fluid. Ethanol was exchanged for PBS and incubated overnight prior to cell seeding.

### Cell isolation

Mice were euthanized and the submandibular glands (SMG) were removed and chopped with a razor blade for 5 min. The tissue was then incubated in Hank’s buffered salt solution (HBSS) containing 15 mM HEPES, 50 U/mL collagenase type II (Thermo Fisher 17101015), and 100 U/mL hyaluronidase (Sigma Aldrich H3506) at 37°C for 30 min. Cells were centrifuged, then resuspended in HBSS with 15 mM HEPES and passed through 100 μm and 20 μm mesh filters to isolate clusters between 20-100 μm. The digestion protocol produces cell cluster sizes evenly distributed between 20 to 100 μm [33]. The isolated clusters were combined with hydrogel precursor solution and seeded in MB array-based chips as described below.

### Cell seeding with hydrogel precursor solution

Isolated submandibular gland cell clusters (20-100 μm) were encapsulated with poly(ethylene glycol) (PEG) hydrogels within MB arrays as previously described [17]. Briefly, the cells were resuspended in hydrogel precursor solution containing 2 mM norbornene-functionalized 4-arm 20 kDa PEG-amine macromers, 4 mM of the dicysteine functionalized MMP degradable peptide (GKKCGPQG↓IWGQCKKG), 0.05 wt% of the photoinitiator lithium phenyl-2,4,6-trimethylbenzoylphosphinate (LAP) [34], and 0.1 mg/mL laminin in PBS [4,5,17,35]. The cell/gel precursor solution (25 μL for 48 well; 20 μL for 96 well) was pipetted onto the MB chips and incubated for 30 min, pipetting every 10 min to redisperse cells that had settled onto the surface of the chip. The hydrogels were polymerized *in situ* using a Hand-Foot 1000 A broad spectrum UV light (UVA: 5 mW/cm^2^; UVB: 0.4 mW/cm^2^) with a UVC filter for 1.5 min and cultured with media (0.5 mL for 48 well; 150 μL for 96 well), with media changes every 2 days. Culture medium consisted of Dulbecco’s Modified Eagle medium (DMEM):Ham’s F-12 Nutrient Mixture (1:1) supplemented with 100 U/mL Penicillin and 100 μg/mL Streptomycin, 2 mM Glutamine, 0.5x N2 supplement, 2.6 ng/mL insulin, 2 nM dexamethasone, 20 ng/mL epidermal growth factor (EGF), and 20 ng/mL basic fibroblast growth factor (bFGF).

### Reduced glutathione assay

A reduced glutathione assay was developed for in-chip measurements by adapting the Cellular Glutathione Detection Assay Kit (Cell Signaling Technology #13859). The monochlorobimane reagent was prepared by reconstitution in DMSO per manufacturer directions. For 96 well plates, 10 μL of prepared reagent (1:50 ratio of monochlorobimane (MCB) to Tris assay buffer, per manufacturer instructions) was added to wells containing 100 μL of culture media and incubated for 30 min at 37 °C, washed with PBS, and imaged using an Olympus IX70 microscope with a DAPI filter (Excitation: 358 nm/Emission: 461 nm).

### Cellular senescence assay

A cellular senescence assay was developed by adapting the Cellular Senescence Detection Kit – SPiDER-βGal (Dojindo Molecular Technologies, Inc SG04). Balifomycin A1 and SPiDER-βGal stock solutions were prepared in DMSO per manufacturer directions. The assay was performed by first incubating the chips with Balifomycin A1 (1:1000 dilution in media) for 1 hr at 37 °C. The solution was removed and replaced with 30 μL of media containing Balifomycin A1 (1:1000 dilution) and SPiDER-βGal (1:500 dilution) and incubated for 45 min at 37 °C. Chips were washed twice with media and imaged using a fluorescence microscope with a Texas Red filter (Excitation: 580 nm/Emission: 604 nm).

### Image quantification, graphing, and statistical analysis

For both the reduced glutathione and cellular senescence assays, images were quantified in ImageJ. Regions of interest (ROIs) were created by thresholding the images on the fluorescence signal (localized to the SGm) and mean intensity of each ROI was measured. Data was graphed and statistical analyses (ANOVA with Tukey’s post-hoc test) were performed using GraphPad Prism 9. Diagrams for Figure 5.2A and Figure 5.4A were created using Biorender (biorender.com).

**Figure 1:**
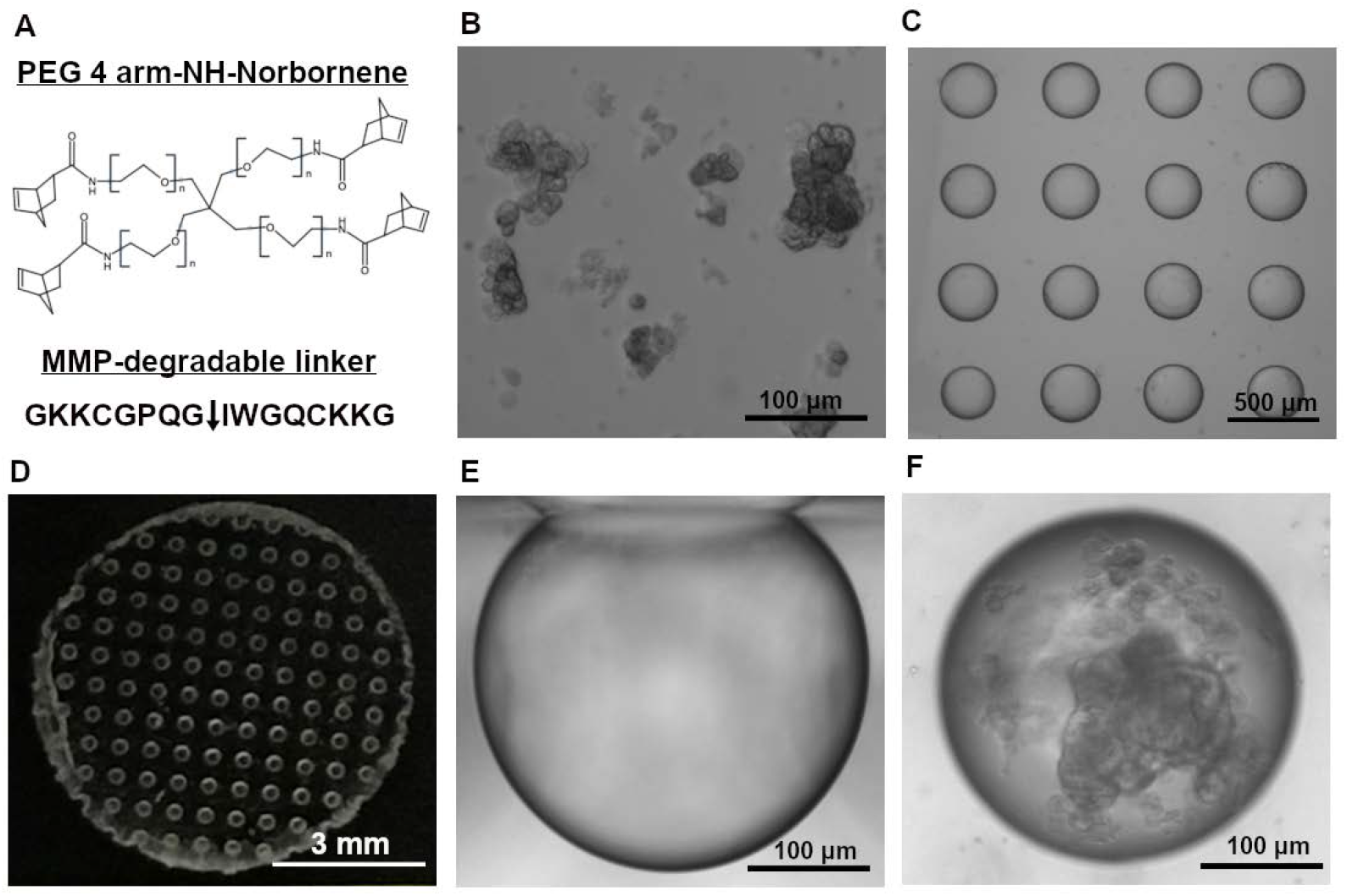
Overview of the components of the salivary gland tissue chip. The tissue chip contains PEG-Norbornene with an MMP-degradable crosslinker (A) and primary salivary gland cell clusters 20-100 μm (B) that are seeded into the MB arrays (C). Macroscale image of an MB chip prior to gluing into a well plate (D). Cross-sectional view of an MB (E). Clusters aggregate to form SGm within the MBs, shown at day 7 (F).

**Figure 2:**
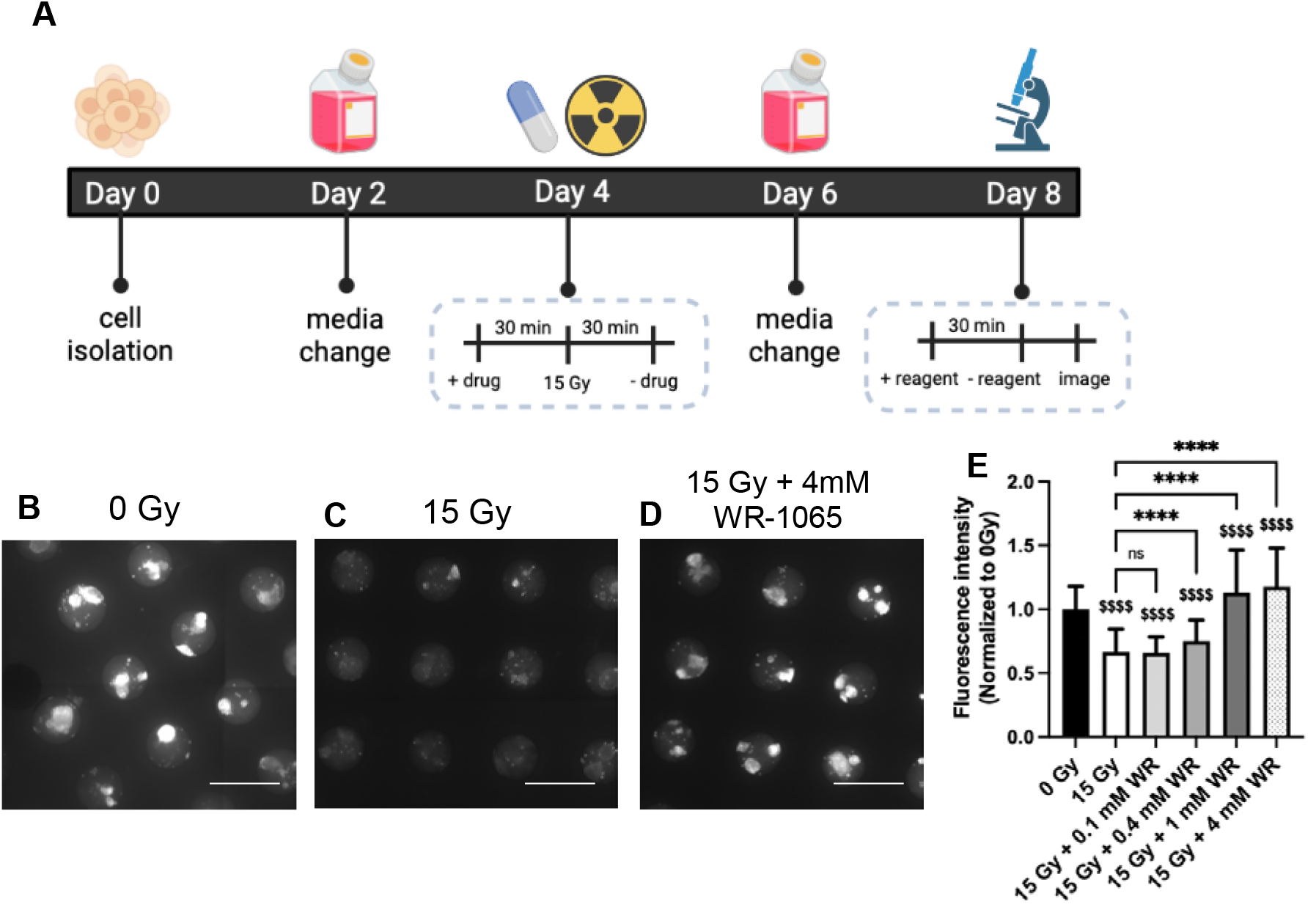
Reduced glutathione assay showed a dose-response to radioprotective drug, WR-1065. Timeline of drug treatment and analysis of glutathione at 4 days post-radiation created using Biorender.com (A). Representative images of the glutathione assay for 0 Gy (B), 15 Gy (C), and 15 Gy + 4 mM WR-1065 (D). Quantification of the fluorescence intensity of individual MBs normalized to 0 Gy (E). Brackets with astericks were compared to 15 Gy: ns = nonsignificant, ****p < 0.0001; Money signs were compared to 0 Gy: $$$$ p < 0.0001; WR = WR-1065; N (# of chips) ≥ 3, n (# of MBs) ≥ 150. Scale bar is 600 μm.

**Figure 3:**
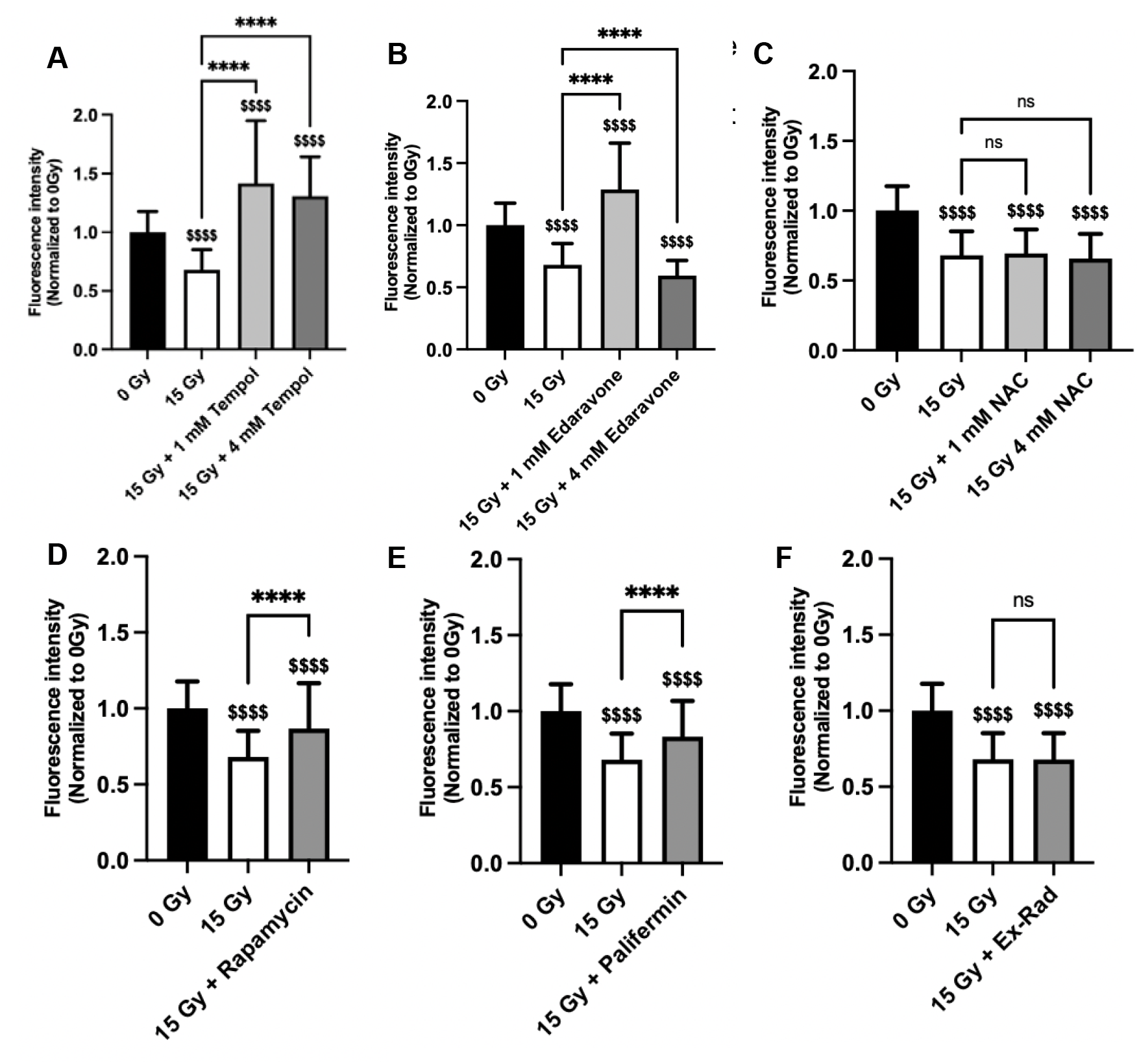
Drugs with reported radioprotective ability are assessed using the reduced glutathione assay. SGm were treated on day 4 for 30 min prior to and 30 min after radiation. Glutathione was analyzed on day 8 for Tempol (A), Edaravone (B), N-acetylcysteine (C), 50 μM Rapamycin (D), 100 ng/mL Palifermin (E), and 50 μM Ex-Rad (F). Brackets with astericks were compared to 15 Gy: ns = nonsignificant, ****p < 0.0001; Money signs were compared to 0 Gy: $$$$ p < 0.0001; NAC = N-acetylcysteine; N (# of chips) ≥ 3, n (# of MBs) ≥ 150.

**Figure 4:**
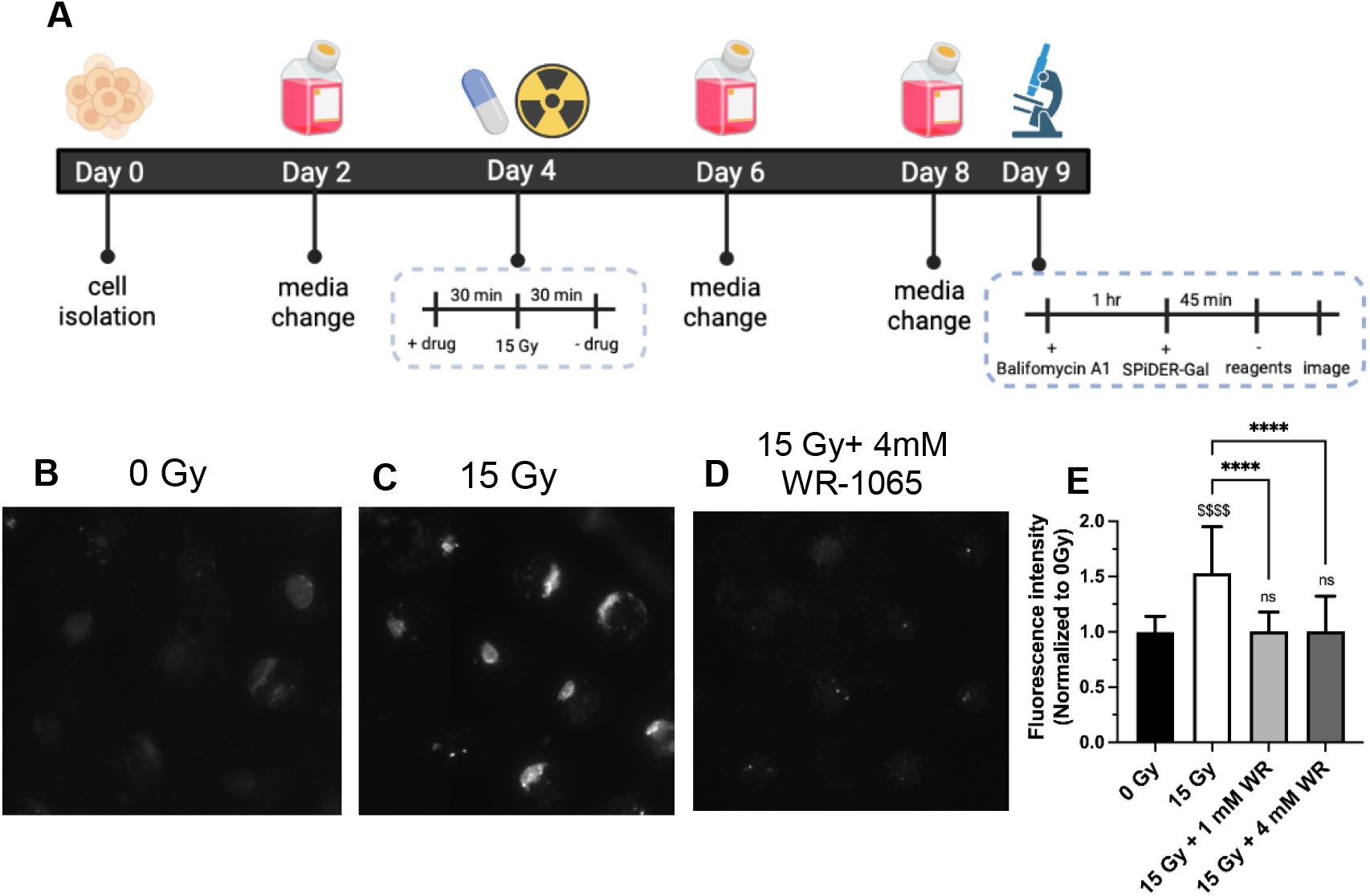
Cellular senescence is increased with radiation and restored with WR-1065 treatment. Timeline of drug treatment and analysis of senescence at 5 days post-radiation created using Biorender.com (A). Representative images of the senescence assay for 0 Gy (B), 15 Gy (C), and 15 Gy + 4 mM WR-1065 (D). Quantification of the fluorescence intensity of individual MBs normalized to 0 Gy (E). Brackets with astericks were compared to 15 Gy: ****p < 0.0001; Money signs were compared to 0 Gy: $$$$ p < 0.0001, ns = nonsignificant; WR = WR-1065; N (# of chips) ≥ 3, n (# of MBs) ≥ 120.

**Figure 5:**
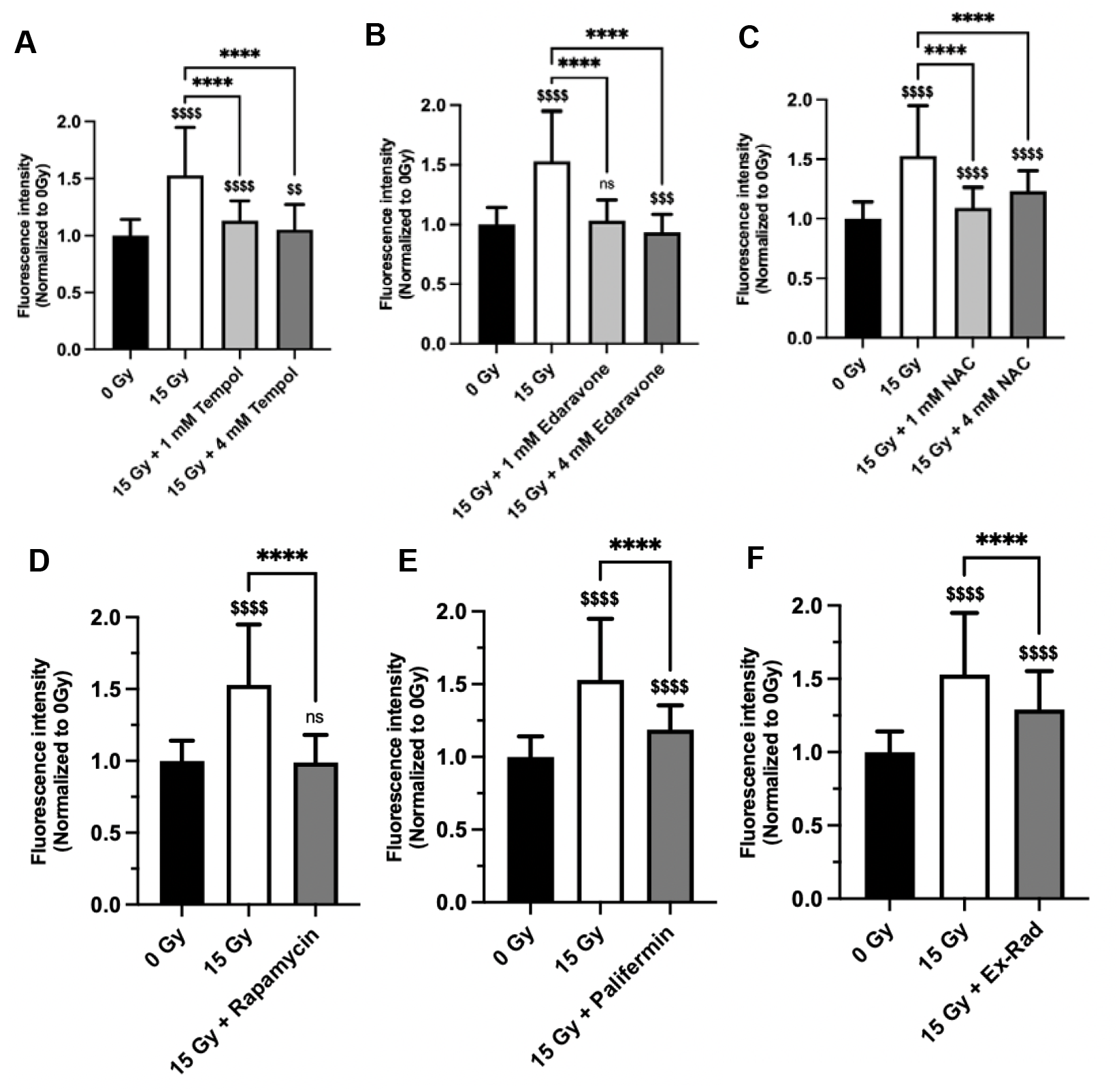
Drugs with reported radioprotective ability were assessed using the senescence assay. SGm were treated on day 5 for 30 min prior to and 30 min after radiation and senescence was analyzed on day 9 for Tempol (A), Edaravone (B), N-acetylcysteine (C), 50 μM Rapamycin (D), 100 ng/mL Palifermin (E), and 50 μM Ex-Rad (F). Brackets with asterisks were compared to 15 Gy: ****p < 0.0001; Money signs were compared to 0 Gy: $$$$ p < 0.0001, $$$ p < 0.001, $$ p < 0.01, ns = nonsignificant; NAC = N-acetylcysteine; N (# of chips) ≥ 3, n (# of MBs) ≥ 120.

### Drug treatment and irradiation

For radioprotection experiments, SGm were cultured in MB-hydrogel chips for 4 days, then drugs were added to the chips 30 min prior to radiation and washed out with media 30 min post-radiation. A dose of 15 Gy ionizing radiation was delivered using a JL Shepherd ^137^Cs irradiator. Drug treatment scheme and radiation dose were established in our previous work [17]. The glutathione and senescence assays were performed at 4 and 5 days post-radiation, respectively. For assay validation experiments, at least 3 chips (N = 3) were used for each drug (Table 1), corresponding to > 100 MBs (n > 100); these values are listed in the figure captions for each experiment. Mean, standard deviation, and statistics were calculated based on the number of MBs (n).

For screening of the Selleck Chemicals compound library, the same treatment scheme was used, with compounds administered at 100 μM. One MB chip (N = 1) was used per compound, with ~40-50 MBs per chip (n = 40-50); statistics were calculated using the number of MBs (n) and compared to 0 Gy controls. Compounds were first screened using the glutathione assay; compounds that were statistically nonsignificant compared to the 0 Gy control (hits) with the glutathione assay were then tested with the senescence assay.

## 3. Results and Discussion

### Overview of the chip

Our previously validated salivary gland tissue chip [17] was used to facilitate high-content screening. The chip platform consists of an array of near-spherical microbubble (MB) cavities formed in poly(dimethyl) siloxane (PDMS). Each chip, containing ~50 MBs, was glued into wells of a 96-well plate (Figure 1C,D,E). Primary salivary gland cell clusters (Figure 1B), suspended in a poly(ethylene glycol) (PEG) hydrogel precursor solution with an MMP-degradable crosslinker (Figure 1A) and the photoinitiator LAP were applied to the chip allowing the clusters to deposit into the MBs. *In situ* polymerization of the hydrogel was achieved using UV light. Over time, the cell clusters aggregate and grow to form SGm (Figure 5.1F).

### Reduced glutathione assay

The reduced glutathione assay was tested at a variety of time points post-radiation to determine the optimal time point for detecting a change between 0 Gy and 15 Gy. Based on the data, the greatest signal separation was measured at 4 days post-irradiation (Figure S1). This timepoint is similar to previous reports on decreases in reduced glutathione content post-radiation [18] and was used for all experiments moving forward.

Next, WR-1065 was tested to determine the effective concentration(s) of WR1065 that prevent changes in reduced glutathione levels post-radiation. SGm in the chip were cultured for 4 days to allow spheres to form [17]. The chip was then treated with WR-1065 for 30 min prior to and during radiation, with the drug washed out with media 30 min post-radiation (Figure 5.2A). This dosing scheme is consistent with amifostine’s use clinically [14] and our previous work [17]. The glutathione assay was performed at 4 days post-radiation (Figure 2A). Example images show the high level of reduced glutathione at 0 Gy (Figure 2B) that is decreased by 15 Gy radiation (Figure 2C) and restored with 4 mM WR-1065 (Figure 2D). Quantification shows that doses of 0.1 mM and 0.4 mM WR-1065 were ineffective at preventing radiation damage, while both 1 mM and 4 mM WR-1065 provided significant protection (Figure 2E). The 4 mM dose corresponds with our previous work on DNA damage markers γH2AX and 53BP1 [17] and values from literature [36] and establishes the range of effective concentrations for WR-1065 treatment *in vitro*.

Since WR-1065 is an antioxidant and mediates radioprotective effects through free radical scavenging and induction of superoxide dismutase expression [37], other antioxidants were tested (Tempol, Edaravone, N-acetylcysteine) at 1 and 4 mM. Tempol exhibited strong radioprotection at both concentrations (Figure 3A), an expected result based on data supporting the use of Tempol as a radioprotectant for the salivary gland [20,38,39]. Edaravone showed complete protection at 1 mM, but none at 4 mM (Figure 3B). Edaravone maintained ~43% of glutathione activity at 100 μM (Figure S2A), suggesting that the optimal concentration range for Edaravone might be lower than WR-1065. This is supported by literature, in which typical concentrations range from 100-1000 μM [23,40,41]. For N-acetylcysteine (NAC), no protection

Next, drugs with non-antioxidant protective mechanisms were tested to assess the assay’s potential in screening drugs that target other pathways. Rapamycin is an mTOR inhibitor that has been reported to restore salivary flow rate post-irradiation in swine [24]. Ex-Rad reduces p53-dependent and independent apoptosis [25]. Palifermin (keratinocyte growth factor) has been reported to stimulate salivary gland stem/progenitor cell expansion post-radiation [27]. Using drug concentrations based on literature [26,28], our data shows partial radioprotection with 50 μM rapamycin (58%, Figure 3D) and 100 ng/mL Palifermin (47%, Figure 3E), with no effect from 50 μM Ex-Rad (Figure 3F).

### Cellular senescence assay

A dosing protocol similar to the glutathione assay was followed to determine the optimal time point post-radiation for detecting a change in senescence, as measured by senescence-associated β-galactosidase activity, between 0 Gy and 15 Gy. Cells were treated with WR-1065 for 30 min prior to and during radiation, with the drug was washed out 30 min post-radiation (Figure 4A). A timepoint of 5 days post-radiation was found to be optimal (Figure S3), similar to literature [19]. An increase in senescence was detected for SGm exposed to 15 Gy (Figure 4C) compared to 0 Gy (Figure 5.4B) that was restored with the addition of WR-1065 (Figure 4D). Quantification shows that both 1 mM and 4 mM WR-1065 treatment resulted in complete radioprotection (Figure 4E), which corroborates results from the reduced glutathione assay.

Next, the senescence assay was tested with the other known radioprotective drugs for validation. Tempol (Figure 5A) and Edaravone (Figure 5B) showed protection at both 1 mM and 4 mM (75% and 91% for Tempol, 94% and 113% for Edaravone, at 1 and 4 mM, respectively). Edaravone also showed 96% protection at 100 μM (Figure S4A). For NAC, 83% and 57% protection was observed with 1 mM and 4 mM (Figure 5C) and 82% at 10 mM (Figure S4B). For non-antioxidant drugs, rapamycin showed complete protection (Figure 5D), while Palifermin provided 64% protection (Figure 5E) and Ex-Rad had minimal impact on radioprotection (45%, Figure 5F).

In general, results for the glutathione and senescence assays were similar (Table S2). Slight differences may be a result of the different drug mechanisms of action and different assay targets. The glutathione assay may be more suitable for detecting antioxidant function (NAC, Tempol, Edaravone), while the senescence assay may be more appropriate for drugs like rapamycin, which has been shown to have anti-senescence properties [42]. These results highlight the trade-offs in developing screening assays. The assays represent indirect markers for DNA damage, a fundamental outcome of radiation-induced cell damage, but were validated for screening a drug library with enhanced throughput compared to immunohistochemical staining for γH2AX. Moreover, the tissue chip array format provides high-content analysis with multiple replicates (40-50) per test.

### Selleck Chemicals library screening

The glutathione and senescence assays were utilized for screening a library of FDA-approved compounds (Selleck Chemicals) at 100 μM. Compounds were first screened using the glutathione assay according to the timeline shown in Figure 2A. Any compound that was statistically nonsignificant with the 0 Gy control was considered a hit (Figure 6, orange circles). Hits with the glutathione assay were then tested with the senescence assay and considered a “double hit” if they were nonsignificant compared to 0 Gy (Figure 6, blue circles). A list of the compounds tested in the drug library and relevant statistics are shown in Appendix B. Overall, 438 compounds from the library were tested with a hit rate of 5.7%, for a total of 25 double hits (Table S3, Table 2). While this hit rate is higher than many drug screening applications (0.1 – 0.3%) [43,44], it is likely because compounds are already FDA-approved, indicating that they have already passed rigorous tests for efficacy, specificity, and toxicity. Additionally, phenotypic screens tend to have a higher hit rate than target-based screens (> 1%) [45,46].

**Figure 6:**
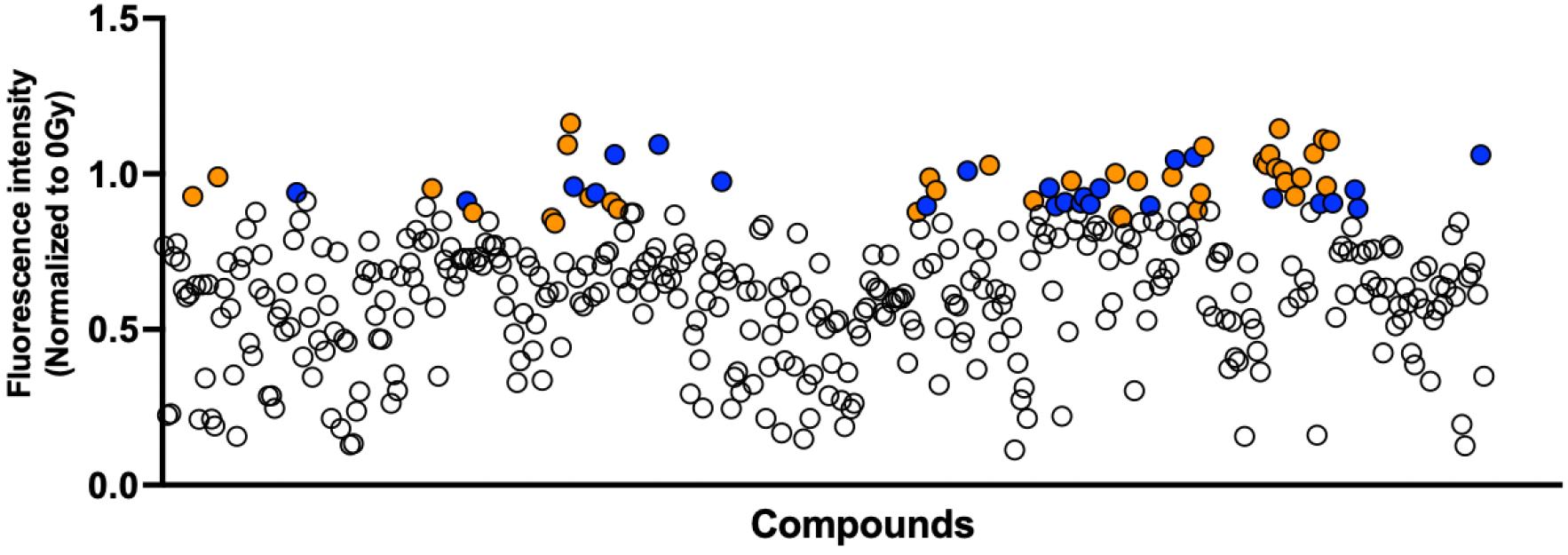
The drug screen identified 25 compounds that protect against post-radiation changes in glutathione and senescence. Each circle represents the normalized fluorescence intensity for the glutathione assay for each compound. White circles are compounds that were not hits. Orange circles are compounds that were hits with the glutathione assay only. Blue circles are compounds that were hits with both assays.

### Drug downselection process

A downselection process was used to triage the 25 double hits for further optimization. Since drugs in the library are FDA-approved, considerable information on their pharmacology in mice and humans is readily available through resources such as PubChem (https://pubchem.ncbi.nlm.nih.gov). Within PubChem, the BioAssay database was created by the National Institute of Health (NIH) as an open repository containing results of small molecule and small interfering RNAs (siRNAs) screening data [47]. BioAssay results were used to identify drug promiscuity to reduce the number of drug candidates to carry forward. Promiscuity refers to a drug’s ability to bind to multiple molecular targets with distinct pharmacological outcomes, often causing unwanted side effects [48]. Thus, drugs exhibiting bioactivity in a large number of assays were deprioritized. Data for each of the double hits was obtained from the database and promiscuity was calculated as the percent of assays reported as “active” (Table 3). Drugs with a high percent (> 10%) were excluded from further testing. Additionally, etidronate, melatonin, and albendazole were excluded due to poor bioavailability [49–51] and eplerenone was excluded due to batch-to-batch variability concerns.

**Table 3:** Drug activity based on data from the BioAssay database. Drugs highlighted in dark gray were excluded from further analysis due to bioactivity in > 10 % of assays. Drugs in light gray were excluded based on poor bioavailability and batch variability (eplerenone).

| Drug | Total Assays | Active (%) |
| --- | --- | --- |
| Glipizide (Glucotrol) | 3141 | 0.54 |
| Etidronate (Didronel) | 1402 | 1.07 |
| Meropenem | 7031 | 1.31 |
| Oxcarbazepine | 2774 | 1.33 |
| Doripenem | 2434 | 1.4 |
| Phenylbutazone | 5565 | 1.55 |
| Enoxacin (Penetrex) | 2673 | 1.87 |
| Cilostazol | 3093 | 2.46 |
| Triamcinolone (Aristocort) | 2345 | 2.73 |
| Didanosine (Videx) | 2840 | 5.67 |
| Eplerenone | 610 | 5.9 |
| Prazosin HCl | 1264 | 6.33 |
| Indomethacin | 10138 | 6.56 |
| Diethylstilbestrol (Stilbestrol) | 5048 | 7.25 |
| Melatonin | 4488 | 8.2 |
| Emtricitabine (Emtriva) | 516 | 9.3 |
| Albendazole (Albenza) | 3086 | 9.3 |
| Rifampin | 8071 | 11.03 |
| Nifedipine (Adalat) | 6190 | 11.47 |
| Rifabutin (Mycobutin) | 1123 | 12.02 |
| Calcitriol (Rocaltrol) | 1786 | 15.23 |
| Lomustine (CeeNU) | 5832 | 16.08 |
| Adefovir Dipivoxil (Preveon) | 929 | 22.71 |
| Curcumin | 3404 | 29.67 |
| Gemcitabine (Gemzar) | 2190 | 46.67 |

For the remaining 13 compounds, the drugs were tested for radioprotection at lower concentrations (1-100 μM) using the glutathione assay. Many of the compounds were only effective at 100 μM and excluded due to a lack of potency. This left seven drugs, phenylbutazone, meropenem, diethylstilbestrol, prazosin HCl, enoxacin, glipizide, and doripenem hydrate, for which a larger range of concentrations was tested using the glutathione (Figure 7) and senescence (Figure 8) assays. Results differed between the glutathione and senescence assays. For example, while a wide range of concentrations were effective for phenylbutazone (Figure 7A) and meropenem (Figure 7B) with the glutathione assay, the trend was less clear for the senescence assay (Figure 8A,B). Diethylstilbestrol showed protection at 10-100 μM for glutathione (Figure 7C) and 50-100 μM for senescence (Figure 8C). Prazosin HCl showed limited efficacy with the glutathione assay (Figure 7D), despite protection at 0.1-100 μM for the senescence assay (Figure 8D), with the opposite trend for enoxacin (Figure 7E; Figure 8E). Moderate results were observed for glipizide and doripenem hydrate for both assays (Figure 7F,G; Figure 8F,G).

**Figure 7:**
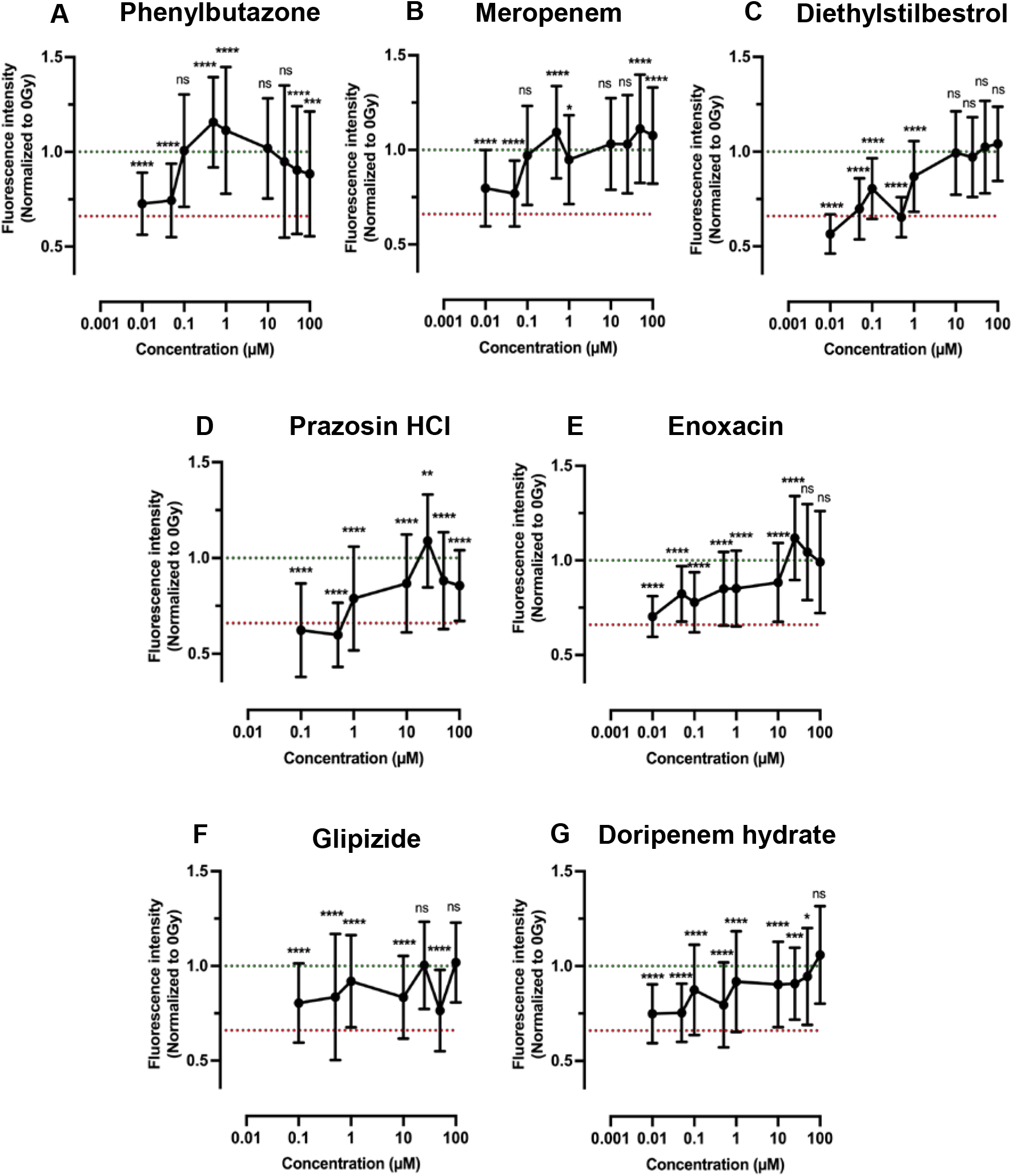
Dose-response data for lead compounds from the drug screen analyzed using the glutathione assay. Results for phenylbutazone (A), meropenem (B), diethylstilbestrol (C), prazosin HCl (D), enoxacin (E), glipizide (F), and doripenem hydrate (G). Green and red lines represent 0 Gy and 15 Gy averages, respectively. Statistics were calculated using ANOVA with Dunnett’s post-hoc test. ns = nonsignificant; **p < 0.01, ***p < 0.001, ****p < 0.0001

**Figure 8:**
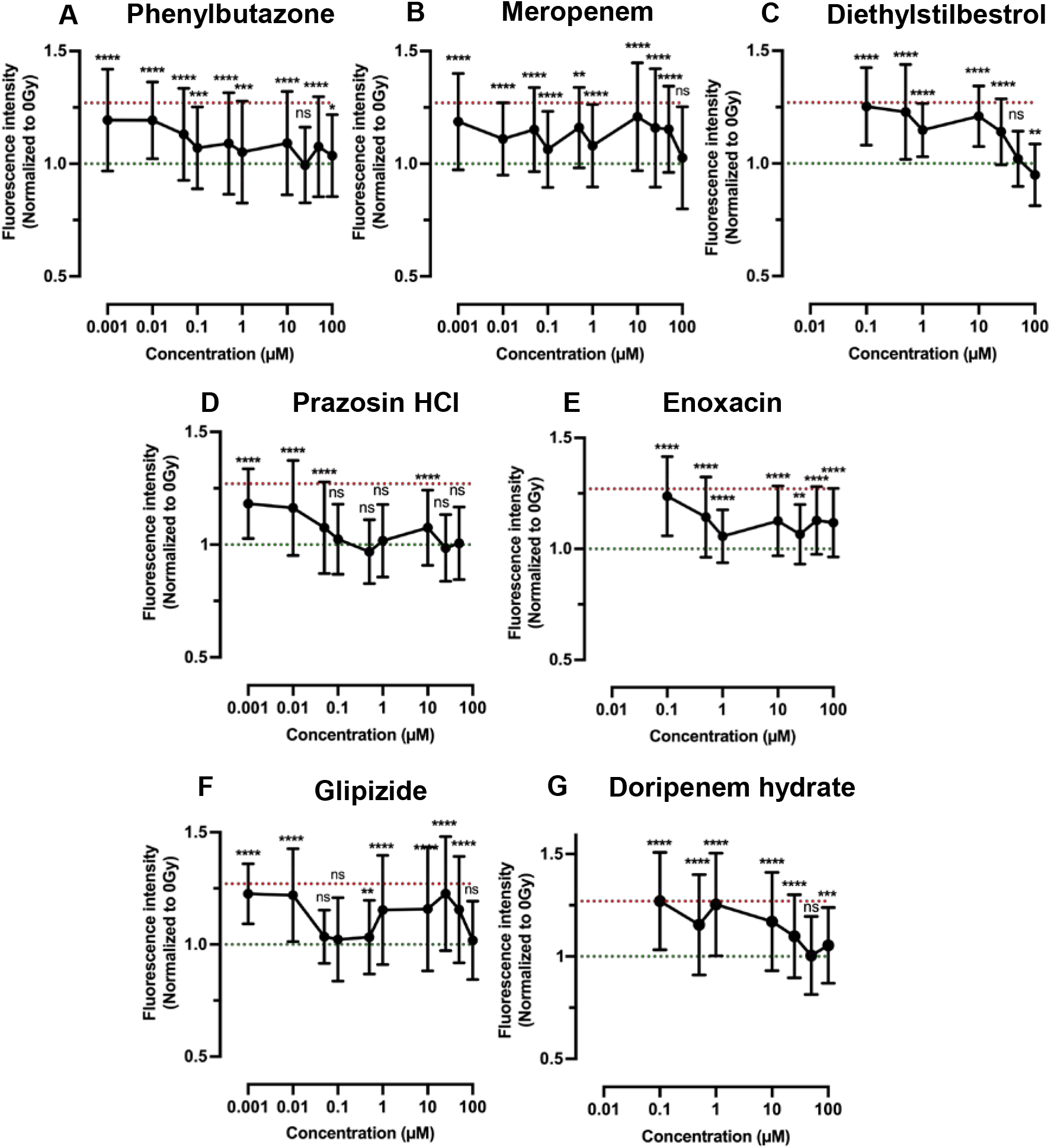
Dose-response data for lead compounds from the drug screen analyzed using the senescence assay. Results for phenylbutazone (A), meropenem (B), diethylstilbestrol (C), prazosin HCl (D), enoxacin (E), glipizide (F), and doripenem hydrate (G). Green and red lines represent 0 Gy and 15 Gy averages, respectively. Statistics were calculated using ANOVA with Dunnett’s post-hoc test. ns = nonsignificant; **p < 0.01, ***p < 0.001, ****p < 0.0001

Estimated EC_50_ values are shown in Table 4. Phenylbutazone showed the most promising results, with low EC_50_ values for both the glutathione (0.083 μM) and senescence (0.048 μM) assays. Phenylbutazone is a non-steroidal anti-inflammatory drug (NSAID) that inhibits cyclooxygenases (COX-1 and COX-2), enzymes that produce prostaglandins [52]. Prostaglandins, specifically PGE_2_ signaling, have been shown to increase in irradiated salivary glands and mitigation of salivary gland damage was achieved through treatment with the anti-inflammatory drug indomethacin [1,53]. Indomethacin also showed radioprotection in our drug screen but was not effective at concentrations lower than 100 μM. Since indomethacin only blocks COX-1, this may explain why phenylbutazone was more effective.

**Table 4:** Estimated EC_50_ values for the top radioprotective drugs. Values were estimated using a nonlinear fit in Prism. “Undetermined” indicates that the software was unable to accurately fit a curve to the data.

| Compound | Estimated EC<sub>50</sub> (μM) | |
| --- | --- | --- |
|  | Glutathione | Senescence |
| Phenylbutazone | 0.083 | 0.048 |
| Meropenem | 0.098 | undetermined |
| Diethylstilbestrol | 0.66 | 37.66 |
| Prazosin HCl | 0.98 | 0.04 |
| Enoxacin | 2.44 | 0.48 |
| Glipizide | 4.03 | undetermined |
| Doripenem hydrate | 8.62 | 16.42 |

Phenylbutazone was originally used for chronic pain for conditions such as arthritis, but has since been restricted to treatment of ankylosing spondylitis due to induction of rare but severe blood disorders, including anemia and leukopenia [52]. However, doses ranged from 300-1000 mg, generating a plasma concentration of 30-50 μg/mL. In contrast, the EC_50_ of 0.083 μM for protecting against radiation-related glutathione changes established in this study equates to a concentration of 26 ng/mL. Thus, the risk of severe adverse effects may be greatly diminished in this context. Additionally, phenylbutazone has excellent bioavailability (up to 90%) [52,54] and a long half-life (50-105 hrs) [52], which may allow for smaller doses, further decreasing risks.

Enoxacin, an antibacterial agent used for treating urinary tract infections [55], has previously been identified as a radioprotector. Using a high-throughput screening method with viability of lymphocytes as the primary readout, two classes of antibiotics (tetracyclines and fluoroquinolones) were identified as robust radioprotectors, including enoxacin [56]. This observation matches up with our drug screening results, in which several antibiotics were identified as double hits, including enoxacin, meropenem, doripenem hydrate, rifampin, and rifabutin. The EC_50_ of 2.4 μM for the glutathione assay is similar to the 13 μM EC_50_ reported for viability of lymphocyte cells [56]. Notably, five other fluoroquinolones reported as radioprotectors (levofloxacin, gatifloxacin, ofloxacin, moxifloxacin, and norfloxacin) [56] were not hits in our drug screen (see Appendix A). These disparities may be related to differences in cell type (salivary gland vs. lymphocyte) or readouts (glutathione/senescence vs. viability).

Glipizide is a sulfonylurea used to treat type 2 diabetes by promoting insulin secretion [57]. It binds to sulfonylurea receptor type 1 (SUR1), which closes ATP-sensitive potassium channels [57]. The buildup of intracellular K^+^ causes membrane depolarization, opening voltage-gated calcium channels [57]. Glyburide and glicliazide, similar sulfonylureas used as anti-diabetic medication, have been described as radioprotectors. Glicliazide was shown to have antioxidant activity [58], while glyburide was suggested to regulate apoptosis by controlling intracellular calcium and the mitochondrial permeability transition (MPT) pore [59]. A patent exists for the use of glyburide as a radioprotective agent (US8883852B2) [59].

## 4. Conclusions

A reduced glutathione assay and a cellular senescence assay were shown to detect radiation damage to salivary gland mimetics (SGm) in a high-content cell culture platform consisting of MMP-degradable PEG hydrogels and microbubble (MB) array technology. The assays were validated with known radioprotective drugs, including WR-1065, the only currently approved preventative therapy for radiation-induced xerostomia, as well as Tempol, Edaravone, N-acetylcysteine, Palifermin, Ex-Rad, and rapamycin. A library of FDA-approved drugs was screened, identifying 25 compounds with radioprotective ability. This list was narrowed down top 3 candidates using the PubChem database and dose-response studies to based on EC_50_ values. Studies are on-going to test these top drug candidates in mice.

## 5. Supplemental Materials

**Figure S1:**
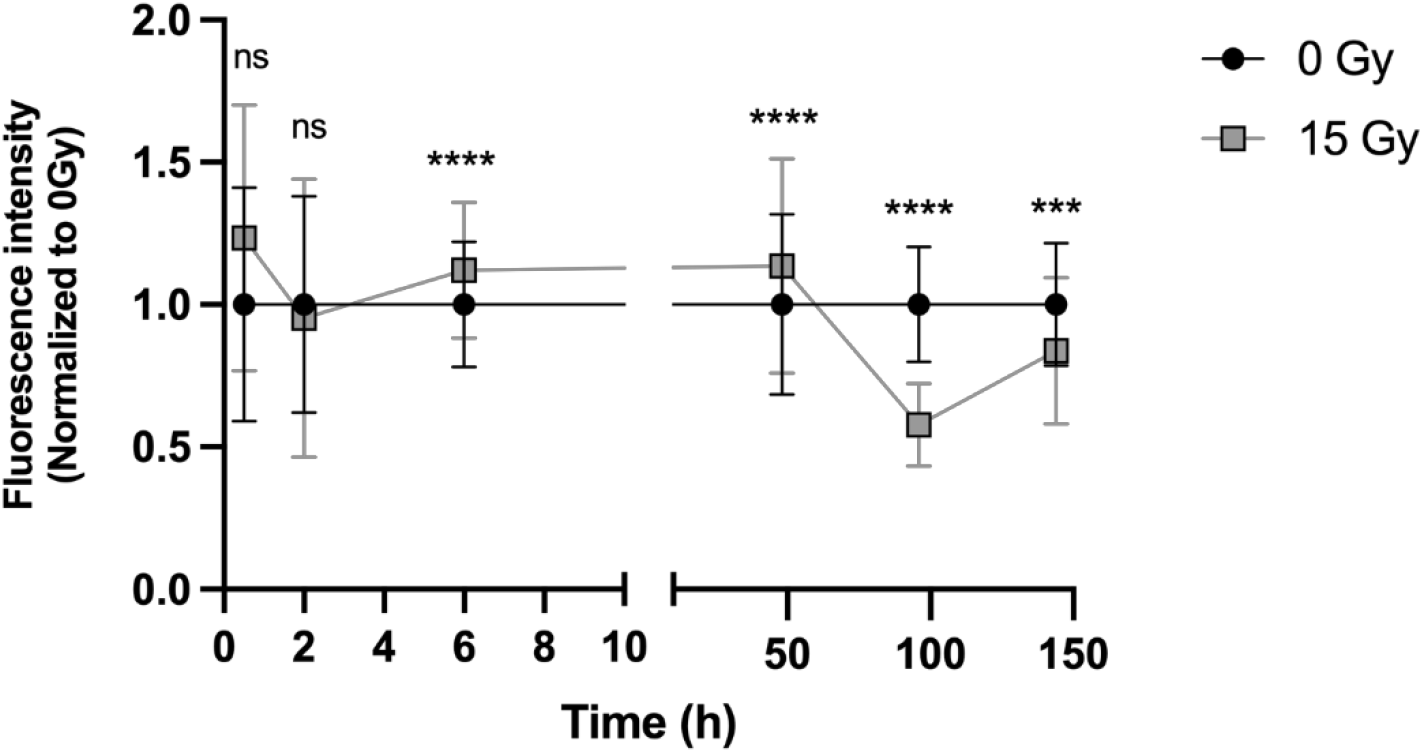
The reduced glutathione assay shows the best signal separation at 4 days post-radiation. Fluorescence intensity is normalized to 0 Gy. Statistics are comparing 0 Gy to 15 Gy at each timepoint. ns = nonsignificant, ****p < 0.0001, ***p < 0.001, N (# of chips) ≥ 3, n (# of MBs) ≥ 120.

**Figure S2:**
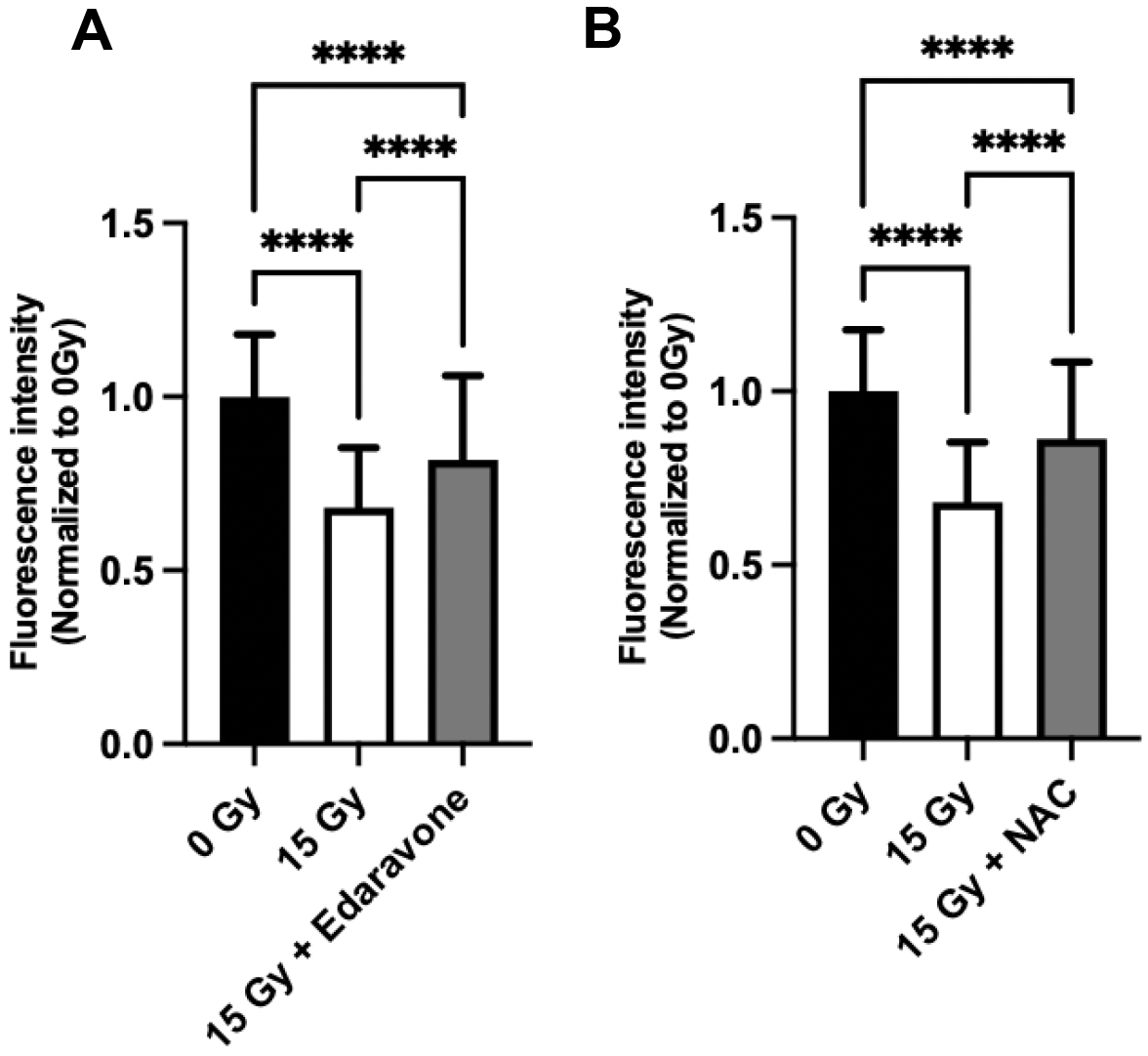
Edaravone and N-acetylcysteine tested at additional concentrations show partial radioprotection. Edaravone (A) tested at 100 μM and N-acetylcysteine (B) tested at 10 mM. ****p < 0.0001; NAC = N-acetylcysteine; N (# of chips) ≥ 3, n (# of MBs) ≥ 200.

**Figure S3:**
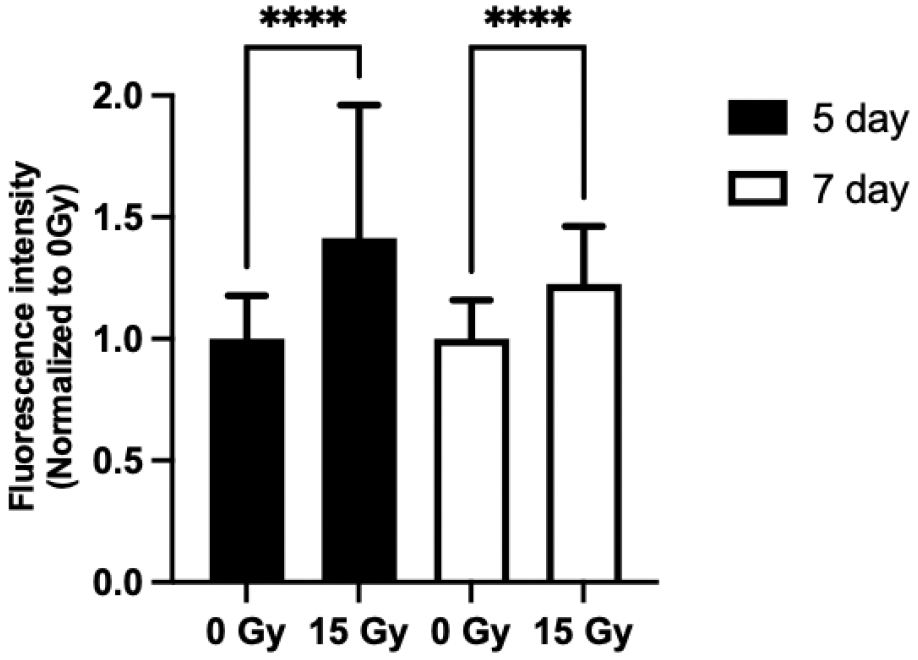
Cellular senescence assay shows the best signal separation at 5 days post radiation. Senescence assay tested at 5 and 7 days. N ****p < 0.0001; N (# of chips) ≥ 4, n (# of MBs) ≥ 180.

**Figure S4:**
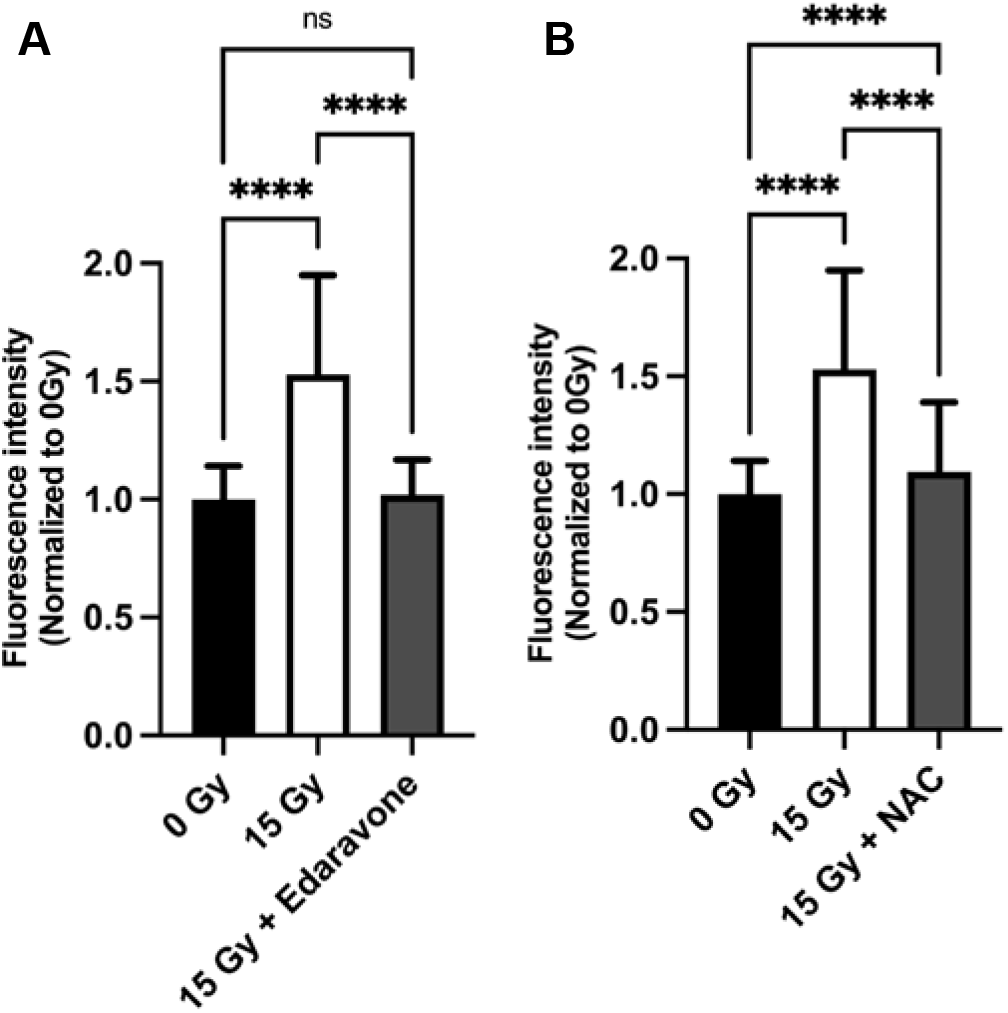
Edaravone and N-acetylcysteine tested at additional concentrations for senescence show complete and partial radioprotection, respectively. Edaravone (A) tested at 100 μM and N-acetylcysteine (B) tested at 10 mM. ****p < 0.0001; NAC = N-acetylcysteine; N (# of chips) = 3, n (# of MBs) ≥ 120.

**Table S1:** Assays tested for detecting radioprotection in the salivary gland tissue chip. Rows in gray are the assays capable of detecting a reliable change between 0 Gy and 15 Gy at the timepoints in bold.

| Assay | Category | Target | Timepoints |
| --- | --- | --- | --- |
| Image-iT Lipid peroxidation | Free radical damage | Lipid peroxidation | 1 - 5 days |
| CellEvent caspase 3/7 | Apoptosis | Caspase 3/7 | 2 - 7 days |
| CellROX | Free radicals | ROS | 1 - 3 days |
| Calcein AM | Cell viability | Esterase activity | 0.5 - 2 hr, 2 - 4 days |
| PrestoBlue | Metabolism | NAD<sup>+</sup>/NADH | 0.5 - 6 hr, 2 - 4 days |
| CellTiterFluor | Metabolism | Aminopeptidase | 0.5 - 6 hr, 2 - 4 days |
| Carbonic anhydrase assay kit | Secretion | Carbonic anhydrase | 2 hr, 2 - 7 days |
| Calcium signaling | Secretion | Calcium signaling response to agonists | 24 hr |
| PAS-AB stain | Secretion | Mucins | 2 - 7 days |
| H<sub>2</sub>DCFDA (2',7'-dichlorodihydrofluorescein diacetate) | Free radicals | ROS | 1 - 8 hr, 1 - 4 days |
| Cellular glutathione detection kit | Antioxidant | Reduced glutathione | 0.5 - 6 hr, 2 - **4 days** |
| Cellular Senescence Detection Kit - SPIDER-βGal | Cellular senescence | SA-β-galactosidase | **5 days**, 7 days |

**Table S2:**
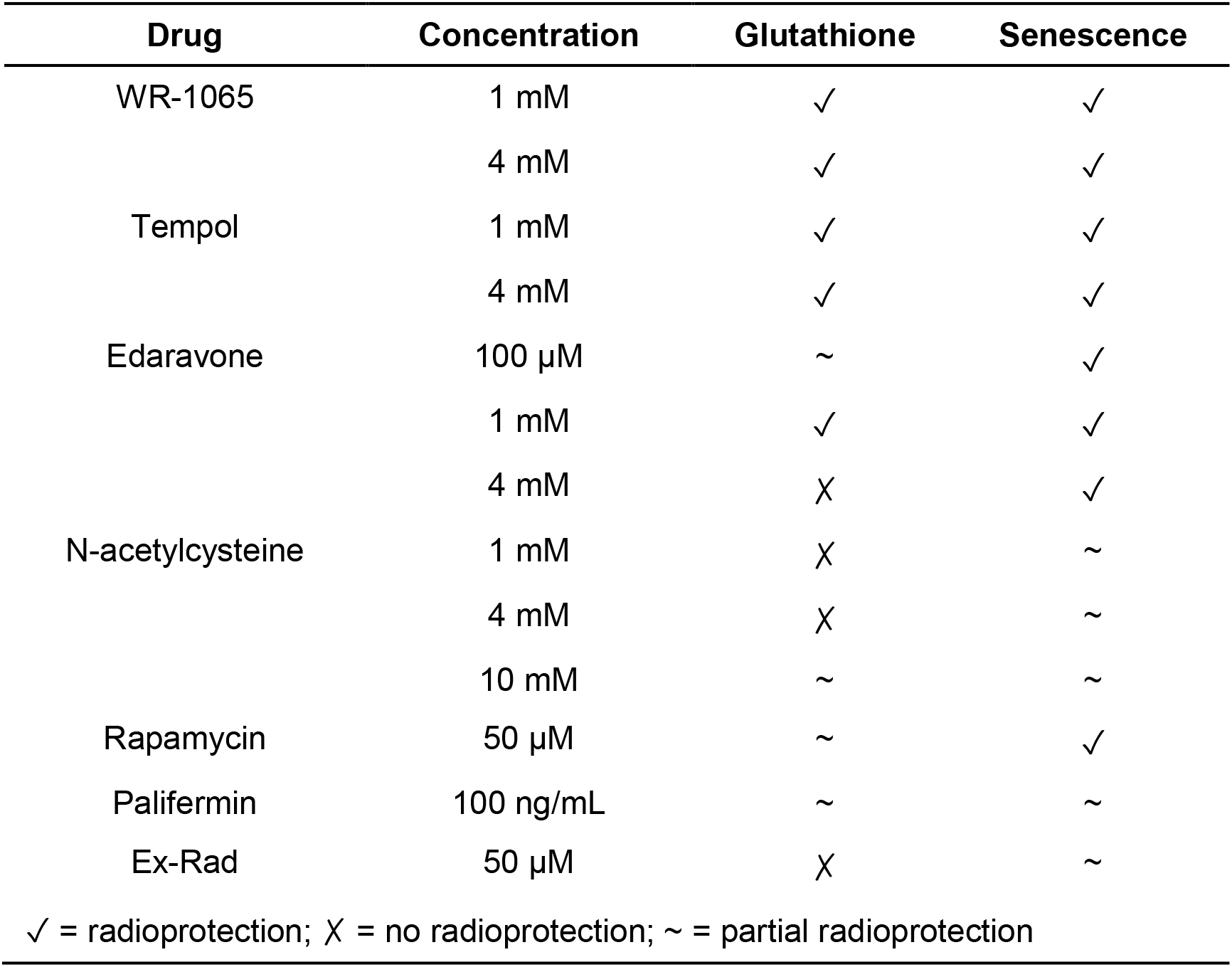
Summary of the results of known radioprotective drugs tested with the glutathione and senescence assays.

| Drug | Concentration | Glutathione | Senescence |
| --- | --- | --- | --- |
| WR-1065 | 1 mM | ✓ | ✓ |
|  | 4 mM | ✓ | ✓ |
| Tempol | 1 mM | ✓ | ✓ |
|  | 4 mM | ✓ | ✓ |
| Edaravone | 100 µM | ~ | ✓ |
|  | 1 mM | ✓ | ✓ |
|  | 4 mM | X | ✓ |
| N-acetylcysteine | 1 mM | X | ~ |
|  | 4 mM | X | ~ |
|  | 10 mM | ~ | ~ |
| Rapamycin | 50 µM | ~ | ✓ |
| Palifermin | 100 ng/mL | ~ | ~ |
| Ex-Rad | 50 µM | X | ~ |

✓ = radioprotection; X = no radioprotection; ~ = partial radioprotection

**Table S3:** Summary of hit rates for the Selleck Chemicals library screening.

| Selleck Chemicals library screening |  |
| --- | --- |
| Drugs screened with glutathione assay | 438 |
| Hits with glutathione assay | 62 |
| Hit rate (percentage) | 14.2% |
| Double hits | 25 |
| Double hit rate (percentage) | 5.7% |

